# Storage stability of non-encapsulated pneumococci in saliva is dependent on null-capsule clade, with strains carrying *aliC* and *aliD* showing a competitive disadvantage during culture enrichment

**DOI:** 10.1101/2025.04.12.648276

**Authors:** Claire S. Laxton, Orchid M. Allicock, Chikondi Peno, Tzu-Yi Lin, Alidia Koelewijn, Femke L. Toekiran, Luna Aguilar, Anna York, Anne L. Wyllie

## Abstract

**Background:** Non-encapsulated *Streptococcus pneumoniae* (NESp) represent up to 19% of circulating pneumococci and exhibit high rates of antimicrobial resistance. Saliva is increasingly used as a pneumococcal carriage study specimen, and we recently developed a qPCR assay to enhance carriage surveillance and characterization of NESpn in saliva. Previous work has established that pneumococci remain viable in unsupplemented saliva for extended periods under various conditions, however these findings may not be applicable to NESp. Therefore, to ensure the robustness of NESp detection in saliva-based carriage studies we evaluated the impact of transport and storage conditions of saliva samples on NESp detection.

**Methods:** Six NESp strains from two clinically relevant NESp null capsule clades (NCC), NCC1 (carrying *pspK*) and NCC2 (carrying *aliC* and *aliD*), were spiked into *lytA*-negative saliva and incubated through various temperatures and freeze-thaw conditions. Endpoints were processed using either culture-enrichment and DNA extraction (CE-DNA), or an extraction-free method without CE, before testing for *lytA* using qPCR. Detection stability was assessed using regression modelling over temperature, time and freeze-thaws.

**Results:** Following CE-DNA, detection of NESp remained stable for ≤24 or ≤72 hours when stored at room temperature or 4°C, respectively, and over 2 freeze-thaw cycles (-80°C), with glycerol-supplementation providing slight benefits. Stability of detection when using CE-DNA depended on NCC; detection of NCC2 strains was lower, and less stable than NCC1. Compared to CE-DNA, extraction-free detection was more stable, with no significant loss over 72 hours at room temperature and over 3 freeze-thaw cycles. With extraction-free detection, there were also negligible differences in detection between NCC1 and NCC2. Additionally, extraction-free detection of NCC1, and less so NCC2, increased over the first 24 hours when stored at 20-30°C, suggesting growth in saliva. Testing of *ΔaliCaliD* and *ΔpspK* mutants revealed these genes increased *in vitro* viability of NCC2 and NCC1, respectively, but did not significantly alter competitive fitness during CE.

**Conclusion:** NCC1 NESp strains exhibit similar stability patterns in unsupplemented saliva as encapsulated pneumococci. NCC2 strains, however, are less resilient during CE, likely due to competition with other oral microbes. Therefore, recovery of NCC2 NESp may be impacted by transport and storage conditions, leading to an underestimation of carriage prevalence when tested using CE-based methods. For the reliable carriage surveillance of NESp, samples should be stored at 4°C soon after collection and at -80°C within 72 hours. Methods which directly detect DNA without CE may provide a less biased accounting of NCC2 strains.

## Introduction

The upper respiratory tract commensal bacterium, *Streptococcus pneumoniae* (pneumococcus) remains the leading global cause of morbidity and mortality from lower respiratory infections. The introduction of pneumococcal conjugate vaccines (PCVs) has led to a substantial reduction in pneumococcal disease caused by up to 21 of the 100+ known pneumococcal serotypes [1, 2]. Concomitantly, PCV use has driven shifts in pneumococcal population dynamics, leading to increased prevalence of non-vaccine targeted serotypes (NVTs) and non-encapsulated strains [3–5].

Non-encapsulated *Streptococcus pneumoniae* (NESp) account for 3-19% of circulating pneumococcal strains and exhibit high rates of multi-drug resistance [6]. NESp are divided into Group I, which retain non-functional capsular genes, and Group II, which completely lack capsular genes in the *cps* locus [7]. Group II NESp are further classified into Null Capsule Clades (NCC) based on the presence of specific genetic elements within the cps locus, including *pspK* (NCC1), *aliC* and *aliD* (NCC2), only *aliD* (historically NCC3) or transposable elements alone (NCC4) [6, 8–10]. While the absence of capsule generally reduces virulence, NCC1 and NCC2 strains have been associated with conjunctivitis outbreaks, otitis media and occasional cases of invasive pneumococcal diseases [6, 11–15].

The genes present in NCC1 and NCC2 strains, *pspK, aliC* and *aliD,* are virulence factors which can compensate for the loss of capsule. AliC and AliD regulate the expression of choline-binding protein AC (CbpAC), which helps reduce C3b deposition on the bacterial surface, thereby enhancing resistance to classical complement-mediated clearance [16, 17]. PspK increases biofilm formation and bacterial adherence to epithelial cells. In turn, this leads to enhanced transmission during influenza co-infection and reduces nasopharyngeal clearance due to interactions with secretory IgA [8, 18, 19].

Carriage prevalence of NESp is increasing, which is concerning due to their heightened propensity for genetic exchange, including antibiotic resistance elements [14, 18, 20–25]. Often, pneumococcal carriage studies neglect to further classify serologically non-typable strains, thus the prevalence of NCC1 and NCC2 NESp is most likely underreported [5, 6, 23, 26]. We recently sought to remedy this by developing a multiplex-qPCR method to simplify the classification NCC1 and NCC2 and increase the inclusion of NESp monitoring in future carriage studies [27].

While nasopharyngeal swabs remain the gold-standard sample for pneumococcal carriage detection [28], swab collection is invasive and resource-intensive, requiring specialized materials and trained healthcare professionals. Saliva is a more accessible sample type than swabs [29] and is increasingly being used for pneumococcal carriage detection [30–35]. As such, the use of saliva for community-based studies continues to be validated and optimized. For example, we and others have demonstrated the high storage stability of respiratory pathogens, including encapsulated pneumococci, in unsupplemented saliva samples [36–39]. However, the stability of NESp, particularly the potentially more virulent NCC1 and NCC2 strains, has not yet been similarly investigated. Therefore, we examined the stability of the detection of NESp in saliva, over different temperatures, timespans and freeze-thaw cycles, to better inform the experimental design of future saliva-based pneumococcal carriage studies and improve the accuracy of NESp carriage estimates.

## Methods

### Saliva samples

Participants were asked to provide whole-mouth unstimulated saliva by passively drooling saliva into a sterile 25 mL polypropylene tube at least 30 min after eating, drinking, or brushing their teeth. Samples were transferred at room temperature to the laboratory within 30 minutes of collection for temporary storage at 4°C, and aliquots were made and stored at -80°C within 12 h.

Each sample was thoroughly screened for absence of pneumococci by incubating an aliquot of each sample at 37°C, and tested at 24, 48 and 72 h for the genes encoding the major pneumococcal autolysin LytA (*lytA)* [40] and pneumococcal iron uptake ABC transporter lipoprotein PiaB (*piaB)* [41, 42], using methods described previously [27]. Saliva samples which tested negative for both *piaB* and *lytA* were thawed, pooled, aliquoted and either placed on ice for immediate use, or stored until needed at -80°C.

### Bacterial isolates

Pneumococcal isolates (**Table 1**) representing the major null-capsule clades (NCC1, 2a and 2b) were obtained from Lance Keller and Larry McDaniel (University of Mississippi Medical Centre, USA), Moon H Nahm (University of Alabama, USA) and Ron Dagan (Ben-Gurion University, Israel) [15, 43, 44].

**Table 1:** Strains of Non-encapsulated Streptococcus pneumoniae used in this study. MLST: multi-locus sequence type, GII-NE Group II Non-Encapsulated, NCC: Null capsule clade,

| Strain | NCC, MLST | Source | Whole Genome Sequence Accession |
| --- | --- | --- | --- |
| **MNZ41** | NCC2b, 6153 | Carriage isolate, South Korea [44] | ASJQ000000000 [45] |
| **MNZ11** | NCC1, 6151 | Carriage isolate, South Korea [44] | ASJW000000000 [45] |
| **MNZ85** | NCC2a, 2315 | Carriage isolate, South Korea [44] | ASJF000030000 [45] |
| **C144.66** | NCC1, 9570 | Adenoiditis isolate, USA [15] | LSMB000000000.1 [15] |
| **D37** | NCC2, NF | Carriage isolate, Israel [43] | JBILPC000000000 [27] |
| **D48** | NCC2, NF | Carriage isolate, Israel [43] | JBILPD000000000 [27] |
| **MNZ1131** | NCC1, 6151 | MNZ11 Δ*pspK* mutant [44] | - |
| **JLB01** | NCC2b, 6153 | MNZ41 Δ*aliCaliD* mutant [17] | - |
| NCC = Null capsule clade, MLST = multi-locus sequence type, NF = MLST not found |  |  |  |

Isolates were plated as a lawn onto tryptic soy agar II (TSA II) plus 5% (v/v) defibrinated sheep blood (blood plates) and incubated at 37°C with 5% CO_2_ overnight. The lawn was harvested into 1 mL brain heart infusion (BHI) medium using a cotton swab and used to inoculate 45 mL BHI medium. Cultures were grown at 37°C with 5% CO_2_ to an optical density at 620 nm (OD_620_) of ∼0.6 absorbance units. Cultures were harvested by centrifugation at 4,000 × *g*, and pellets were resuspended in 5 to 10 mL BHI medium supplemented with 10% (v/v) glycerol and stored at −80°C. The bacterial concentration (CFU/mL) of each sample were determined by colony counting of serially diluted samples cultured on blood plates and incubated at 37°C with 5% CO_2_ overnight. Preliminary work from this study (data not shown), and other studies have found that due to aggregate formation, a phenomenon not observed with encapsulated pneumococcus, vigorous vortexing is required between serial dilution of NESp [46]. The recommended approach, to dilute into individual 1.5 mL microcentrifuge tubes with vortexing for 5-10 secs between dilutions, was adopted here to ensure accurate colony counting.

### Experimental design

For each isolate listed in **Table 1**, concentrated stocks were diluted in BHI then spiked into saliva at final concentrations of 10^3^ and 10^4^ CFU/mL (final concentration of BHI <3% v/v), as described previously [36]. Spiked saliva samples were incubated at 4°C, room temperature (∼20°C), and 30°C. At 24 h, 48 h, and 72 h, the samples were vortexed, and 100 μL was removed from each for culture enrichment (plated immediately) to test for strain viability as described previously [27, 36]. A further 50 µL of each sample was aliquoted and stored at -80°C, for extraction-free pneumococcal detection [39].

To determine the effect of freeze-thawing on the stability of pneumococcal detection, saliva was spiked with pneumococci to final concentrations of 10^3^ and 10^4^ CFU/mL as above (unsupplemented) or supplemented with BHI with 50% glycerol (final concentration 30% BHI v/v and 15% glycerol v/v). Aliquots were stored for a minimum of 3 h at either −20°C or −80°C to allow complete freezing, before being thawed at room temperature. Once thawed, the samples were vortexed, and 100 μL was removed from each sample for culture enrichment before the remainder was returned to −20°C or −80°C. This process was repeated twice more. Where sample volume was insufficient for processing due to technical issues, the datapoint was excluded.

To further examine the recovery of strains immediately following spiking, representative NCC1 (MNZ11) and NCC2 (MNZ41) strains were spiked at 15,000 CFU/mL into five different *lytA* and *piaB* negative saliva samples collected and prepared as above, in a matched pairs fashion. Stains were spiked into 1 mL BHI in the same way in triplicate. Following spiking, samples were vortexed for 10 s then plated for culture enrichment within 10 mins.

### Saliva sample processing by CE-DNA-extraction or extraction-free methods

Culture-enriched saliva samples were thawed at room temperature and DNA was extracted from 200 μL of each sample using the MagMAX Viral/Pathogen Nucleic Acid Isolation Kit (MVP I) using a KingFisher Apex instrument (ThermoFisher Scientific), with modifications [35]. The method is referred to here as CE-DNA-extraction.

Spiked saliva samples were additionally processed using an extraction-free method, without culture enrichment, as described previously [39]. This method involves a lysis step with proteinase K, heat inactivation at 95°C for 10 minutes and immediate transfer of the treated sample (which we will refer to as lysate) for pneumococcal detection via qPCR.

### Detection of pneumococcal carriage

Each sample (2.5 µL of either DNA template or extraction-free lysate) was tested using the same dualplex qPCR targeting *piaB* and *lytA* as described above. Genomic DNA extracted from a pneumococcus serotype 19A strain (*lytA* and *piaB* positive) was included in every plate as a positive control, using at least two standard concentrations (0.0001-1 ng/µL). Assays were run on a CFX96 Touch instrument (Bio-Rad) under the following conditions: 95°C for 3 min followed by 45 cycles of 98°C for 15 s and 60°C for 30 s. Samples were considered positive for pneumococci when the Cq values for both genes were ≤40 and within 2 Cq of each other and negative controls were undetectable. Since NESp strains MNZ41, C144.66 and D48 are naturally *piaB* negative, only *lytA* positivity was considered for samples spiked with these strains [27]. Plate-to-plate variation was corrected for by multiplying each sample Cq by an adjustment factor as described previously [27].

### Statistical analyses

Linear regression using R (version 3.5.2) was conducted to evaluate the impacts of time and temperature (time) or freeze-thaw cycle and supplementation (freeze thaw) on the detection of pneumococci from spiked saliva samples using the *lytA* adjusted Cq. Interaction terms were used to evaluate whether the effects of time and temperature (time), freeze-thaw cycle and supplementation (freeze thaw) and starting concentration (both) varied by strain. The ΔCq value represents the change in the mean Cq value from freshly spiked saliva under each condition (categorical). *P* values of less than 0.05 were considered significant. Tabulated model results are available in the Supplementary data.

To further study viability following CE-DNA-extraction of NCC1 vs NCC2 strains immediately after spiking into saliva or BHI, a mixed-effects analysis was employed using GraphPad Prism (version 10.3.0) with Tukey’s multiple comparisons, utilizing a restricted maximum likelihood algorithm due to different sized groups. The experiment was conducted in a repeated measures fashion and the ΔCq value represents the change in the mean value between each strain. *P* values of less than 0.05 were considered significant.

## Results

### Detection of culture-enriched NESp depends on time, temperature, NCC and storage conditions

When tested following CE-DNA-extraction, the stability of detection of NESp over time was dependent upon temperature (**Figure 1A**). There was no significant loss in detection over 72 h when stored at 4°C (ΔCq= 1.01, p= 0.083, 95% CI: -0.14 - 2.16). However, detection dropped significantly after 48 h when stored at RT (ΔCq= 3.44, p< 0.001, 95% CI: 1.50 - 5.38), and after 24 h when stored at 30°C (ΔCq= 4.86, p< 0.001, 95% CI: 2.76 - 6.97). In addition, stability of detection over time and temperature depended on NCC. When stratified by NCC, detection of NCC2 strains, but not NCC1, dropped significantly after 72 h when stored at 4°C (ΔCq= 1.68, p= 0.035, 95% CI: 0.12 – 3.24), after 48 h when stored at RT (ΔCq= 3.92, p= 0.006, 95% CI: 1.20 - 6.64), and for less than 24 h when stored at 30°C (ΔCq= 7.00, p< 0.001, 95% CI: 4.38 – 9.60).

**Figure 1:**
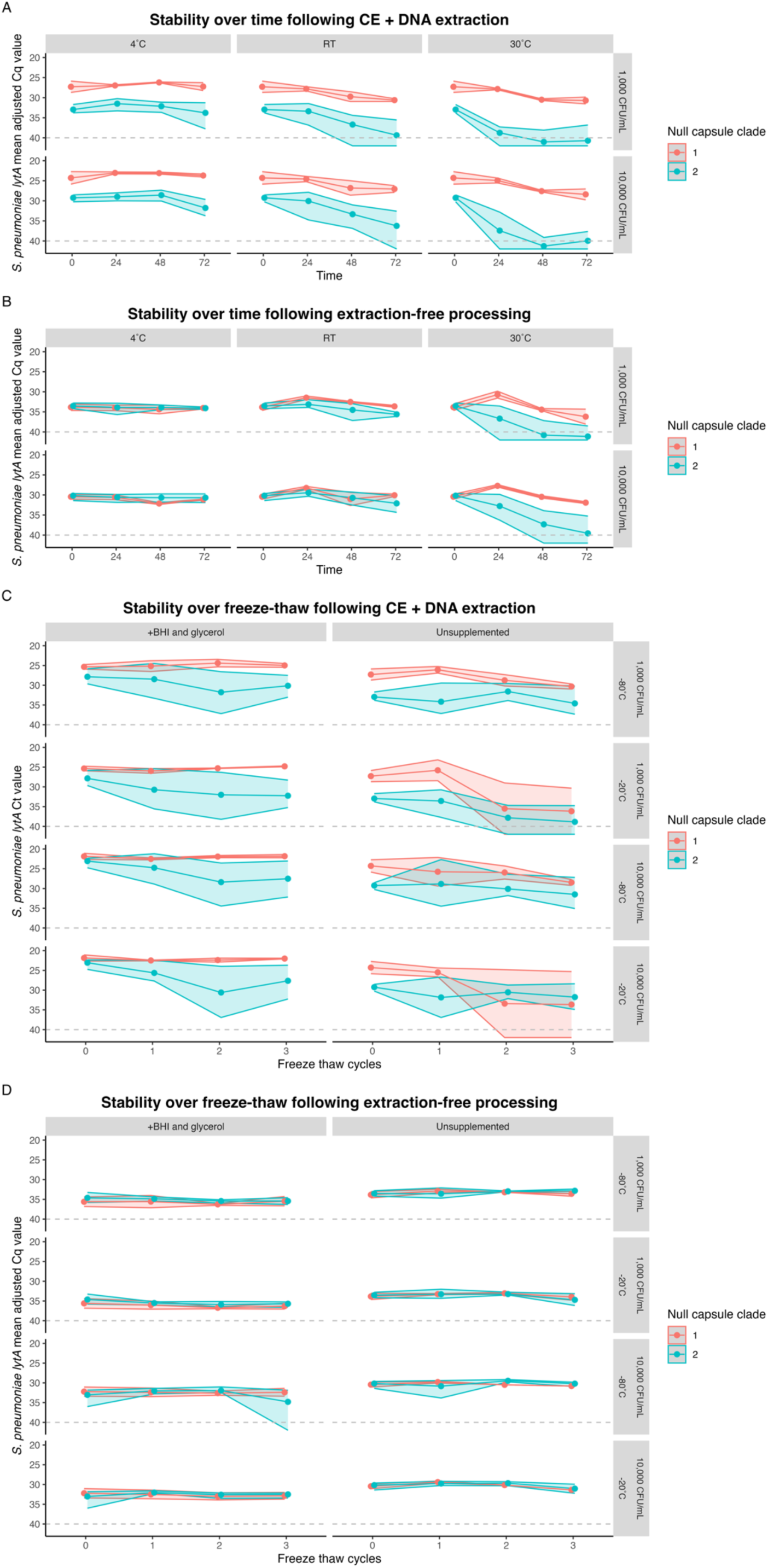
Detection of NCC1 NESp over time, temperature and freeze-thaw is more stable than that of NCC2 when spiked into saliva. Panel A and B show the detection of NESp over time at diFerent temperatures (4°C, RT= room temperature and 30°C) following either culture-enrichment and DNA extraction (CE + DNA) or extraction-free methods, respectively. Saliva was initially spiked at either 1,000 or 10,000 CFU/mL, as indicated by the right-hand label. Panel C and D show the detection of NESp over 3 freeze-thaw cycles following either CE + DNA or extraction-free methods, respectively. Saliva was initially spiked into unsupplemented saliva or saliva supplemented with BHI plus glycerol at either 1,000 or 10,000 CFU/mL and cycled through either -20°C or -80°C, as indicated by the right-hand label. Data are shown as the mean adjusted Cq for lytA for each of NCC1 (n=2, red) or NCC2 (n=4, blue) with the range shown as a shaded ribbon.

We next assessed the effect of freeze-thaw cycles on detection stability of NESp (**Figure 1C**). For samples that were processed by CE-DNA-extraction, detection stability of NESp across freeze-thaw cycles was dependent on storage temperature. There was a significant reduction in detection by the second freeze-thaw cycle for samples stored at -20°C (ΔCq= 4.46, p< 0.001, 95% CI: 2.25 – 6.68) and by the third cycle for samples stored at -80°C (ΔCq= 2.34, p= 0.008, 95% CI: 2.25 – 6.68), suggesting a slightly greater freeze-thaw stability when stored at -80°C.

When tested using CE-DNA-extraction, adding BHI+glycerol to the saliva significantly increased overall bacterial detection following freeze-thaws at -80°C (ΔCq unsupplemented vs glycerol= 3.79, p< 0.001, 95% CI: 2.58 – 5.00) and -20°C (ΔCq unsupplemented vs glycerol= 5.22, p< 0.001, 95% CI: 3.65 - 6.78), however glycerol supplementation did not affect the stability of detection through consecutive freeze-thaws (ΔCq 0 h vs 72 h= 1.62, p= 0.251, 95% CI: -1.16 – 4.41) (**Figure 1C**). Similarly, while overall detection of NCC2 was lower than that of NCC1 strains (ΔCq= 4.20, p< 0.001, 95% CI: 3.17 – 5.25), stability of detection across freeze thaw cycles was unaffected by NCC strain (ΔCq= 0.42, p= 0.781, 95% CI: -2.56 – 3.40).

### Processing of NESp using an extraction-free method leads to greater stability of detection which is less dependent on storage conditions and NCC

To better understand the impact of culture-enrichment on the detection of NESp strains which underwent the same storage conditions, spiked saliva samples were also tested using the extraction-free method, which does not include culture enrichment. Similarly to when tested using CE-DNA-extraction, detection was stable over 72 h when stored at 4°C (ΔCq= 0.50, p= 0.127, 95% CI: -0.15 – 1.15). However, when stored at room temperature there was a slight increase in overall detection at 24 h (ΔCq= -1.14, p= 0.017, 95% CI: -2.07 – -0.21), which became a decrease in detection by 72 h (ΔCq= 1.24, p= 0.10, 95% CI: 0.31 – 2.16), suggesting some strains may have grown a small amount in the saliva. At 30°C, overall detection dropped after 48 h (ΔCq= 4.91, p< 0.001, 95% CI: 2.86 – 6.97).

Unlike with CE-DNA-extraction, there was no difference in stability of detection of NCC at either 4°C or room temperature when processed using the extraction-free method (**Figure 1B**). Detection according to NCC only deviated when stored at 30°C, in which detection of NCC2 strains decreased significantly after 24 h (ΔCq= 2.88, p< 0.027, 95% CI: 0.35 – 5.40), whereas detection of NCC1 strains, like at room temperature, increased at 24 h (ΔCq= -2.93, p< 0.002, 95% CI: -4.55 – 1.30), before decreasing by 72 h (ΔCq= 1.92, p< 0.025, 95% CI: 0.29 – 3.54). This suggests that NCC1 strains, and less so NCC2 strains can grow in whole saliva at temperatures between 20-30°C.

Detection of NESp remained stable over all three freeze-thaw cycles when processed using the extraction free method (ΔCq= 0.47, p= 0.058, 95% CI: -0.02 – 0.95), and we did not observe any impact on NESp detection over repeated freeze-thaw cycle by NCC strain (ΔCq= -0.14, p= 0.470, 95% CI: -0.51 – 0.23), or storage temperature (ΔCq= -0.12, p= 0.507, 95% CI: --0.23 – 0.47) (**Figure 1D**). Unexpectedly, addition of glycerol supplementation resulted in slightly lower overall detection when samples were processed using the extraction-free method (ΔCq unsupplemented vs supplemented= - 2.22, p< 0.001, 95% CI: 2.57 - 1.87). Glycerol is typically considered a qPCR enhancer and cryoprotectant [47], however it is possible that the addition of glycerol in this concentration or formulation reduced the efficiency of the proteinase K or heat lysis step, resulting in less DNA available for qPCR.

Overall, there were twice as many samples in which *lytA* was not detectable at endpoint following CE-DNA-extraction (7.4%) than extraction-free processing (3.4%) (**Table 2**). Almost all the samples in which *lytA* could not be detected (32/663) were originally spiked with an NCC2 strain, except for four unsupplemented saliva samples which were spiked with a NCC1 strain and underwent ≥2 freeze thaw cycles.

**Table 2:** Percentage of samples from which NESp was not detectable (ND) stratified by treatment group.

| Processing method | Strain | Temperature (% ND) | Freeze thaw (% ND) |  | Total (% ND) |
| --- | --- | --- | --- | --- | --- |
|  |  |  | Unsupplemented | +BHI and glycerol |  |
| CE & DNA extraction | NCC1 | 0/48 (0.0%) | 4/32 (12.5%) | 0/32 (0.0%) | 4/112 (3.6 %) |
|  | NCC2 | 18/96 (18.8%) | 3/64 (4.7%) | 0/64 (0.0%) | 21/224 (9.4%) |
| Extraction-free | NCC1 | 0/48 (0.0%) | 0/28 (0.0%) | 0/32 (0.0%) | 0/108 (0.0%) |
|  | NCC2 | 10/96 (10.4%) | 0/59 (0.0%) | 1/64 (1.6%) | 11/219 (5.0%) |
| Total (% ND) |  | 28/288 (9.7%) | 7/183 (3.8%) | 1/192 (0.5%) | 36/663 (5.4%) |

Following the observation that NCC2 strains were less stable in saliva than NCC1 strains following culture enrichment, we sought to determine if this was driven by the presence of *aliC* and *aliD*. Strains belonging to NCC1 (MNZ11) and NCC2 (MNZ41) were spiked into saliva and BHI in parallel and immediately tested using the CE-DNA-extraction method. Recovery of viable NCC1 strains was significantly higher than NCC2 following spiking into saliva (mean ΔCq= -3.52, 95% CI: -4.67 – -2.37, p<0.0001; **Figure 2A**). However, there was no significant difference when spiked into BHI (mean ΔCq= - 0.06, 95% CI: -1.19 – 1.08, p=0.9972; **Figure 2B**). Recovery of both strains was lower and more varied in saliva than in BHI, suggesting that a component of saliva has an impact on bacterial viability during culture enrichment, which was more pronounced for NCC2 strains.

**Figure 2:**
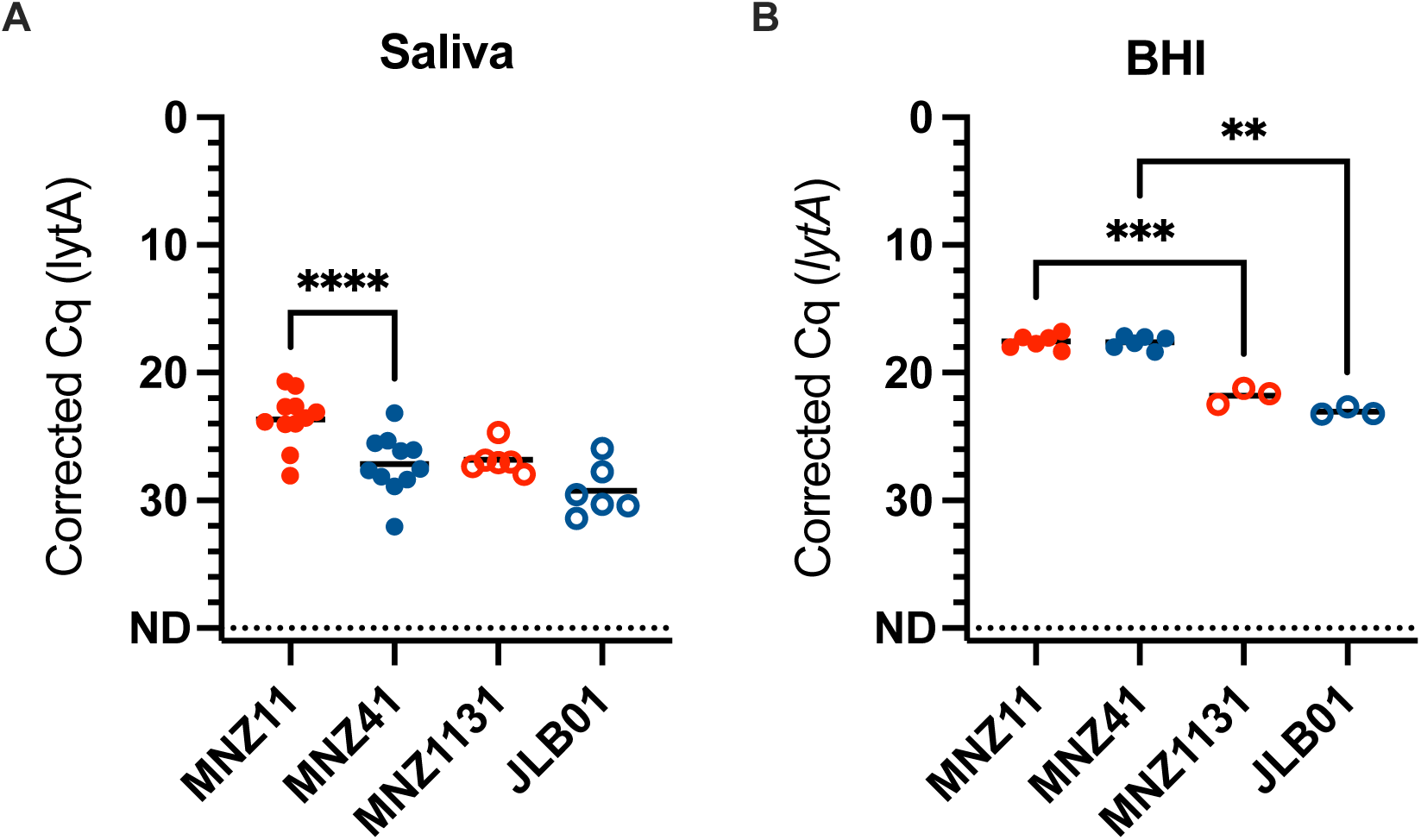
Recovery of NCC2 NESp is lower than NCC1 immediately after spiking into saliva, but not BHI, and this is not dependant on aliC or aliD. MNZ11 (red, filled circles) is NCC1 and MNZ41 (blue, filled circles) is NCC2, MNZ1131 (red, open circles) and JLB01 (blue, open circles) are pspK and aliC/aliD gene knockouts of MNZ11 and MNZ41, respectively. Strains were spiked at a final concentration of 15,000 CFU/mL into either BHI (n=6 for MNZ11 and MNZ41, n=3 for MNZ1131 and JLB01) or matched pairs of lytA and piaB negative saliva from healthy individuals (n=11 for MNZ11 and MNZ41, n=6 for MNZ1131 and JLB01). Line is the mean, significance is shown for multiple comparisons between MNZ11 vs MNZ41, MNZ11 vs MNZ1131 and MNZ41 vs JLB01, **p<0.005, ***p<0.0005, ****p<0.0001.

Knockout mutants MNZ1131 (*ΔpspK*) and JLB01 (*ΔaliCaliD*) were also tested, and while recovery in BHI was significantly lower for both than their wildtypes (**Figure 2B**), there were no significant differences in recovery in saliva. This suggests that while *pspK and aliC* and *aliD* do play a role in overall cell viability, they are not primarily driving the effect on viability of capsule loss during culture enrichment in saliva.

## Discussion

Carriage studies are reporting general increases in the prevalence of NESp, which is of concern particularly in the case of NCC1 and NCC2 strains, due to the increased virulence of these strains, and their propensity to acquire AMR genes [5, 6, 23–25]. There has also been an increase in recent years in the use of saliva as a sample type for investigating pneumococcal carriage, however the impact of saliva sample storage on the detection and recovery of NESp has not yet been described [30, 32, 34, 41, 48–50].

Therefore, in this study we sought to build upon the existing evidence examining encapsulated pneumococci viability in saliva across a range of storage conditions and determine if similar findings applied to NESp. We previously demonstrated that the viability of pneumococci remained stable in self-collected, unsupplemented saliva samples stored at 4°C to 30°C for up to 24 hours, and a portion of viable pneumococci could be detected for up to 72°C, though at lower quantities [36]. Findings from this current study demonstrate that while NCC1 strains exhibit equivalent storage stability in saliva to encapsulated strains [36], NCC2 strains are less able to survive culture enrichment, leading to lower detection than NCC1 following CE-DNA-extraction (**Figure 1A and C**, **Figure 2A**).

Overall, when tested following the extraction-free method, detection was generally lower compared with CE-DNA-extraction, which was expected, due to the lack of enrichment and DNA concentration steps. Interestingly though, compared with CE-DNA-extraction, the extraction-free method led to more stable detection of NESp over both time (**Figure 1B**) and freeze-thaw cycles (**Figure 1D**), with negligible variation between NCC1 and NCC2. There were also fewer endpoint samples with a total loss of detection when using the extraction-free method compared with CE-DNA-extraction (3.4% vs 7.4%, **Table 2**).

Culture enrichment amplifies all viable gentamycin resistant strains in a sample, the totality of which is harvested for detection [30, 41]. Thus, for pneumococcal detection using culture enrichment, any pneumococcal strains present must be both viable at the time of plating, and able to survive on the culture plate during incubation. In the case of saliva, culture enrichment can include numerous mitis-group streptococci, which compete for resources and space [51]. Meanwhile, the extraction-free method used in this study consists simply of heat and enzymatic lysis step and direct-qPCR, meaning bacteria in the sample do not need to be viable to be detected.

Therefore, it was hypothesised that the observed reduction in recovery of NCC2 strains was due to reduced cell survival in saliva and/or growth during culture enrichment. When we compared the recovery of cells immediately after spiking into either saliva or BHI, we found that recovery of NCC2 strains was significantly lower than that of NCC1 strain when culture-enriched form saliva, but not BHI (**Figure 2**). Our findings suggest that while all cells were viable at the time of plating, NCC2 strains, but not NCC1, had a growth disadvantage in saliva during culture-enrichment. Potential explanations for this include a lower competitive fitness of NCC2 against other gentamicin-resistant bacteria present in saliva, or a salivary factor, such as a protease, that specifically inhibits their growth. Further studies are needed which test the recovery of NCC1 and NCC2 strains from either heat-treated or filtered saliva, to systematically determine the salivary component driving reduced NCC2 fitness during CE. Nevertheless, these data suggest that while CE may enhance detection of encapsulated pneumococci and NCC1 NESp, it can cause undue suppression of NCC2 NESp detection and possibly contribute to underreporting of NESp prevalence. Given that the extraction-free method is cheaper and less laborious than CE-DNA-extraction [39], it may be advisable to employ this, or other culture-free methods to ensure accurate detection of NESp.

This study examined stability at two different bacterial loads, 1,000 and 10,000 CFU/mL, and the observed stability trends were generally replicated between each, with the expected ∼3 Cq difference (**Figure 1**). This suggests that the detection and quantification methods employed here were robust. Moreover, the recovery of representative NCC1 and NCC2 strains spiked into BHI was both highly replicable and similar between strains (**Figure 2**), suggesting that the observed variations between strains could not be explained by variations in the quantity or viability of bacteria being spiked into saliva. However, this study was limited by testing the stability of only a small number of NESp strains, and only in the Group II, NCC1 and 2 clades. Given the inter-strain differences observed here, it would be interesting to assess Group I NESp stability, as knowledge of the impact of capsule loss on stability of pneumococci in saliva is essential for informing future carriage study sample collection and storage practices.

NESp prevalence is likely underreported in carriage surveillance due to limited focus on the detection and characterisation of these strains. We recently sought to address this by developing a method to aid the detection of NESp, particularly in oral sample types [27]. To further address the knowledge gap around NESp carriage, this current study aimed to determine the conditions for saliva collection, storage and testing for optimal NESp detection. As such, we have demonstrated that NESp can be reliably detected in saliva for up to 72 hours when stored at 4°C, or 24 hours when stored at room temperature. We have also demonstrated that while glycerol supplementation slightly improves NESp detection when using culture enrichment, it is not otherwise necessary. We also showed that NCC1 strains exhibit comparable resilience to encapsulated pneumococci, while NCC2 strains appear to be at a competitive disadvantage during culture enrichment. Recoverability of viable NCC1 and NCC2 strains was not significantly related the presence of either *pspK* or *aliC*/*aliD* genes, respectively (**Figure 2**). Further investigation is required to understand the mechanism underlying our observation of reduced NCC2 fitness in saliva during CE. These findings underscore the importance of refining detection methodologies and considering strain-specific differences when designing future pneumococcal carriage studies.

## Acknowledgments

Thanks to Lance Keller and Dan Weinberger for their advice, and for kindly providing bacterial strains used in this study, as well as to Larry McDaniel and Mary Marquart for their suggestions on study methodology.

## Author contributions

Conceptualization: CSL, OMA, AY, ALW; Methodology: CSL, OMA, CP, ALW, AY; Investigation: CSL, OMA, CP, AMK, TL, FT, LA, AY; Formal Analysis and Visualization: CSL, CP; Supervision: CSL, AY, ALW; Writing – original draft: CSL, CP, OMA; Writing-review & editing: CSL, OMA, CP, AY, ALW.

## Conflict of interest and Funding disclosures

ALW has received consulting and/or advisory board fees from Pfizer, Merck, Diasorin, PPS Health, Co-Diagnostics, and Global Diagnostic Systems for work unrelated to this project and was the Principal Investigator on research grants from Pfizer, Merck, and NIH RADx UP to Yale University and from NIH RADx, Balvi.io, and Shield T3 to SalivaDirect, Inc. ALW is currently employed by Pfizer, Inc. The authors have no conflicts of interest to declare in relation to this work.

## Ethics Statement

Saliva was collected from healthy adult participants for this study. Potential participants were informed about the purpose and procedure of the study and consented to study participation through the act of providing a saliva sample; the requirement for written informed consent was waived by the Institutional Review Board of the Yale Human Research Protection Program (Protocol ID 2000029374) [36].

## References

1. Manna S, Ortika BD, Werren JP, Pell CL, Gjuroski I, et al. Streptococcus pneumoniae serotype 33H: a novel serotype with frameshift mutations in the acetyltransferase gene wciG. Pneumonia 2025 17:1 2025;17:1–9.

2. Blacklock CB, Weinberger DM, Perniciaro S, Wyllie AL. Streptococcus pneumoniae serotypes. https://pneumococcalcapsules.github.io/serotypes/ (2025, accessed 3 April 2025).

3. Lo SW, Gladstone RA, van Tonder AJ, Lees JA, du Plessis M, et al. Pneumococcal lineages associated with serotype replacement and antibiotic resistance in childhood invasive pneumococcal disease in the post-PCV13 era: an international whole-genome sequencing study. Lancet Infect Dis 2019;19:759–769.

4. Waight PA, Andrews NJ, Ladhani Shamez N and Sheppard CL, Slack MPE, Miller E. Effect of the 13-valent pneumococcal conjugate vaccine on invasive pneumococcal disease in England and Wales 4 years after its introduction: an observational cohort study. Lancet Infect Dis 2015;15:535–543.

5. Nunes S, Félix S, Valente C, Simões AS, Tavares DA, et al. The impact of private use of PCV7 in 2009 and 2010 on serotypes and antimicrobial resistance of Streptococcus pneumoniae carried by young children in Portugal: Comparison with data obtained since 1996 generating a 15-year study prior to PCV13 introduction. Vaccine 2016;34:1648–1656.

6. Keller LE, Robinson DA, McDaniel LS. Nonencapsulated Streptococcus pneumoniae: Emergence and Pathogenesis. mBio;7. Epub ahead of print 22 March 2016. DOI: 10.1128/MBIO.01792-15.

7. Hathaway LJ, Stutzmann Meier P, Bättig P, Aebi S, Mühlemann K. A homologue of aliB is found in the capsule region of nonencapsulated Streptococcus pneumoniae. J Bacteriol 2004;186:3721–3729.

8. Keller LE, Jones C V, Thornton JA, Sanders M E and Swiatlo E, Nahm MH, et al. PspK of Streptococcus pneumoniae increases adherence to epithelial cells and enhances nasopharyngeal colonization. Infect Immun 2013;81:173–181.

9. Thompson CD, Bradshaw JL, Miller WS, Vidal AGJ, Vidal JE, et al. Oligopeptide transporters of nonencapsulated streptococcus pneumoniae regulate CbpAC and PspA expression and reduce complement-mediated clearance. mBio;14.

10. Mohale T, Wolter N, Allam M, Ndlangisa K, Crowther-Gibson P, et al. Genomic analysis of nontypeable pneumococci causing invasive pneumococcal disease in South Africa, 2003–2013. BMC Genomics;17. Epub ahead of print 2016. DOI: 10.1186/S12864-016-2808-X.

11. Park IH, Geno KA, Sherwood LK, Nahm MH, Beall B. Population-Based Analysis of Invasive Nontypeable Pneumococci Reveals That Most Have Defective Capsule Synthesis Genes. PLoS One 2014;9:e97825.

12. Hanage WP, Kaijalainen T, Saukkoriipi A, Rickcord JL, Spratt BG. A Successful, Diverse Disease-Associated Lineage of Nontypeable Pneumococci That Has Lost the Capsular Biosynthesis Locus. J Clin Microbiol 2006;44:743.

13. Xu Q, Kaur R, Casey JR, Sabharwal V, Pelton S, et al. Nontypeable Streptococcus pneumoniae as an Otopathogen. Diagn Microbiol Infect Dis 2011;69:200.

14. Martin CS, Bradshaw JL, Pipkins HR, McDaniel LS. Pulmonary Disease Associated With Nonencapsulated Streptococcus pneumoniae. Open Forum Infect Dis;5. Epub ahead of print 1 July 2018. DOI: 10.1093/OFID/OFY135.

15. Dixit C, Keller LE, Bradshaw JL, Robinson DA, Swiatlo E, et al. Nonencapsulated Streptococcus pneumoniae as a cause of chronic adenoiditis. IDCases 2016;4:56–58.

16. Thompson CD, Bradshaw JL, Miller WS, Vidal AGJ, Vidal JE, et al. Oligopeptide Transporters of Nonencapsulated Streptococcus pneumoniae Regulate CbpAC and PspA Expression and Reduce Complement-Mediated Clearance. mBio;14. Epub ahead of print 28 February 2023. DOI: 10.1128/mbio.03325-22.

17. Bradshaw JL, Pipkins HR, Keller LE, Pendarvis JK, McDaniel LS. Mucosal Infections and Invasive Potential of Nonencapsulated Streptococcus pneumoniae Are Enhanced by Oligopeptide Binding Proteins AliC and AliD. mBio 2018;9:e02097–17.

18. Takeuchi N, Ohkusu M, Wada N, Kurosawa S, Miyabe A, et al. Molecular typing, antibiotic susceptibility, and biofilm production in nonencapsulated Streptococcus pneumoniae isolated from children in Japan. Journal of Infection and Chemotherapy 2019;25:750–757.

19. Sakatani H, Kono M, Nanushaj D, Murakami D, Takeda S, et al. A Novel Pneumococcal Surface Protein K of Nonencapsulated Streptococcus pneumoniae Promotes Transmission among Littermates in an Infant Mouse Model with Influenza A Virus Coinfection. Infect Immun;90. Epub ahead of print 1 February 2022. DOI: 10.1128/IAI.00622-21.

20. Feldman C, Anderson R. Recent advances in the epidemiology and prevention of Streptococcus pneumoniae infections. F1000Res;9. Epub ahead of print 2020. DOI: 10.12688/F1000RESEARCH.22341.1.

21. Bradshaw JL, McDaniel LS. Selective pressure: Rise of the nonencapsulated pneumococcus. PLoS Pathog 2019;15:e1007911.

22. Takeuchi N, Ohkusu M, Hishiki H, Fujii K, Hotta M, et al. First report on multidrug-resistant non-encapsulated Streptococcus pneumoniae isolated from a patient with pneumonia. Journal of Infection and Chemotherapy 2020;26:749– 751.

23. Langereis JD, de Jonge MI. Non-encapsulated Streptococcus pneumoniae, vaccination as a measure to interfere with horizontal gene transfer. Virulence 2017;8:637.

24. Yokota S ichi, Tsukamoto N, Sato T, Ohkoshi Y, Yamamoto S, et al. Serotype replacement and an increase in non-encapsulated isolates among community-acquired infections of Streptococcus pneumoniae during post-vaccine era in Japan. IJID Regions 2023;8:105.

25. Kawaguchiya M, Urushibara N, Aung MS, Kudo K, Ito M, et al. Clonal lineages and antimicrobial resistance of nonencapsulated Streptococcus pneumoniae in the post-pneumococcal conjugate vaccine era in Japan. Int J Infect Dis 2021;105:695–701.

26. Sá-Leao R, Pinto F, Aguiar S, Nunes S, Carriço JA, et al. Analysis of invasiveness of pneumococcal serotypes and clones circulating in portugal before widespread use of conjugate vaccines reveals heterogeneous behavior of clones expressing the same serotype. J Clin Microbiol 2011;49:1369–1375.

27. Laxton CS, Toekiran FL, Lin T-Y, Lomeda BD, Hislop MS, et al. An abundance of aliC and aliD genes were identified in saliva using a novel multiplex qPCR to characterize group II non-encapsulated pneumococci with improved specificity. bioRxiv 2024;2024.11.26.624535.

28. Satzke C, Turner P, Virolainen-Julkunen A, Adrian P V., Antonio M, et al. Standard method for detecting upper respiratory carriage of Streptococcus pneumoniae: Updated recommendations from the World Health Organization Pneumococcal Carriage Working Group. Vaccine 2013;32:165–179.

29. Laxton CS, Peno C, Hahn AM, Allicock OM, Perniciaro S, et al. The potential of saliva as an accessible and sensitive sample type for the detection of respiratory pathogens and host immunity. Lancet Microbe 2023;4:e837–e850.

30. Wyllie AL, Chu MLJN, Schellens MHB, Gastelaars JVE, Jansen MD, et al. Streptococcus pneumoniae in Saliva of Dutch Primary School Children. PLoS One 2014;9:e102045.

31. Wróbel-Pawelczyk I, Ronkiewicz P, Wanke-Rytt M, Rykowska D, Górska-Kot A, et al. Pneumococcal carriage in unvaccinated children at the time of vaccine implementation into the national immunization program in Poland. Scientific Reports 2022 12:1 2022;12:1–11.

32. Wyllie AL, Rümke LW, Arp K, Bosch AATM, Bruin JP, et al. Molecular surveillance on Streptococcus pneumoniae carriage in non-elderly adults; little evidence for pneumococcal circulation independent from the reservoir in children. Scientific Reports 2016 6:1 2016;6:1–9.

33. Almeida ST, Paulo AC, Froes F, De Lencastre H, Sá-Leão R. Dynamics of Pneumococcal Carriage in Adults: A New Look at an Old Paradigm. J Infect Dis 2021;223:1590–1600.

34. Krone CL, Wyllie AL, Van Beek J, Rots NY, Oja AE, et al. Carriage of Streptococcus pneumoniae in Aged Adults with Influenza-Like-Illness. PLoS One 2015;10:e0119875.

35. Wyllie AL, Mbodj S, Thammavongsa DA, Hislop MS, Yolda-Carr D, et al. Persistence of Pneumococcal Carriage among Older Adults in the Community despite COVID-19 Mitigation Measures. Microbiol Spectr;11. Epub ahead of print 15 June 2023. DOI: 10.1128/SPECTRUM.04879-22/SUPPL_FILE/REVIEWER-COMMENTS.PDF.

36. Allicock OM, York A, Waghela P, Yolda-Carr D, Weinberger DM, et al. Impact of Temporary Storage Conditions on the Viability of Streptococcus pneumoniae in Saliva. mSphere;7. Epub ahead of print 21 December 2022. DOI: 10.1128/MSPHERE.00331-22.

37. Marín-Echeverri C, Pérez-Zapata L, Álvarez-Acevedo L, Gutiérrez-Hincapié S, Adams-Parra M, et al. Diagnostic performance, stability, and acceptability of self-collected saliva without additives for SARS-CoV-2 molecular diagnosis. Eur J Clin Microbiol Infect Dis 2024;43:1127–1138.

38. Allicock OM, Lin T-Y, Fajardo KT, Yolda-Carr D, Hislop MS, et al. Exploring the potential of a saliva-based, RNA-extraction-free PCR test for the multiplexed detection of key respiratory pathogens. medRxiv 2023;2023.10.04.23296240.

39. Peno C, Lin T-Y, Hislop MS, Yolda-Carr D, Farjado K, et al. A low-cost culture- and DNA extraction-free method for the molecular detection of pneumococcal carriage in saliva. Microbiol Spectr. Epub ahead of print 19 July 2024. DOI: 10.1128/SPECTRUM.00591-24.

40. Carvalho MDGS, Tondella ML, McCaustland K, Weidlich L, McGee L, et al. Evaluation and improvement of real-time PCR assays targeting lytA, ply, and psaA genes for detection of pneumococcal DNA. J Clin Microbiol 2007;45:2460–2466.

41. Trzciński K, Bogaert D, Wyllie A, Chu MLJN, van der Ende A, et al. Superiority of Trans-Oral over Trans-Nasal Sampling in Detecting Streptococcus pneumoniae Colonization in Adults. PLoS One 2013;8:e60520.

42. Wyllie AL, Pannekoek Y, Bovenkerk S, van Engelsdorp Gastelaars J, Ferwerda B, et al. Sequencing of the variable region of rpsB to discriminate between Streptococcus pneumoniae and other streptococcal species. Open Biol;7. Epub ahead of print 1 September 2017. DOI: 10.1098/RSOB.170074.

43. Tóthpál A, Desobry K, Joshi SS, Wyllie AL, Weinberger DM. Variation of growth characteristics of pneumococcus with environmental conditions. BMC Microbiol;19. Epub ahead of print 26 December 2019. DOI: 10.1186/S12866-019-1671-8.

44. Park IH, Kim KH, Andrade AL, Briles DE, Mcdaniel LS, et al. Nontypeable Pneumococci Can Be Divided into Multiple cps Types, Including One Type Expressing the Novel Gene pspK. mBio;3. Epub ahead of print 2012. DOI: 10.1128/MBIO.00035-12.

45. Keller LE, Thomas JC, Luo X, Nahm MH, McDaniel LS, et al. Draft Genome Sequences of Five Multilocus Sequence Types of Nonencapsulated Streptococcus pneumoniae. Genome Announc 2013;1:e00520–13.

46. Carr MA, Marcelo D, Lovell KM, Benton AH, Tullos NA, et al. Absence of Streptococcus pneumoniae Capsule Increases Bacterial Binding, Persistence, and Inflammation in Corneal Infection. Microorganisms;10. Epub ahead of print 1 April 2022. DOI: 10.3390/MICROORGANISMS10040710.

47. Schaudien D, Baumgärtner W, Herden C. High preservation of DNA standards diluted in 50% glycerol. Diagnostic Molecular Pathology 2007;16:153–157.

48. Rayack EJ, Askari HM, Zirinsky E, Lapidus S, Sheikha H, et al. Routine saliva testing for SARS-CoV-2 in children: Methods for partnering with community childcare centers. Front Public Health 2023;11:1003158.

49. Waghela P, Davis R, Campbell M, Datta R, Hislop MS, et al. Detection of pneumococcal carriage in asymptomatic healthcare workers. medRxiv 2024;2024.07.19.24309369.

50. Almeida ST, Pedro T, Paulo AC, de Lencastre H, Sá-Leão R. Re-evaluation of Streptococcus pneumoniae carriage in Portuguese elderly by qPCR increases carriage estimates and unveils an expanded pool of serotypes. Scientific Reports 2020 10:1 2020;10:1–7.

51. Okahashi N, Nakata M, Kuwata H, Kawabata S. Oral mitis group streptococci: A silent majority in our oral cavity. Microbiol Immunol 2022;66:539–551.

