## Supplementary data for "Storage stability of non-encapsulated pneumococci in saliva is dependent on null-capsule clade, with strains carrying *aliC* and *aliD* showing a competitive disadvantage during culture enrichment"

**CE + DNA extraction stability over temperature and time**

| Full model | Reference | Estimate ΔCq | P-value | Lower CI | Upper CI |
|---|---|---|---|---|---|
| TemperatureRT | 4C | -0.50 | 0.647 | -2.65 | 1.65 |
| Temperature30 | 4C | -2.72 | 0.014 | -4.87 | -0.57 |
| Time24 | Time0 | -0.84 | 0.346 | -2.60 | 0.92 |
| Time48 | Time0 | -0.88 | 0.324 | -2.64 | 0.88 |
| Time72 | Time0 | 1.01 | 0.256 | -0.74 | 2.77 |
| Concentration10000 | Concentration1000 | -2.75 | <0.001 | -3.47 | -2.03 |
| NCC_2 | NCC_1 | 5.90 | <0.001 | 4.58 | 7.21 |
| **Interactions** | | | | | |
| TemperatureRT:Time24 | Time0, 4C | 1.38 | 0.273 | -1.10 | 3.87 |
| Temperature30:Time24 | Time0, 4C | 5.70 | <0.001 | 3.22 | 8.19 |
| TemperatureRT:Time48 | Time0, 4C | 4.32 | 0.001 | 1.83 | 6.80 |
| Temperature30:Time48 | Time0, 4C | 8.66 | <0.001 | 6.18 | 11.15 |
| TemperatureRT:Time72 | Time0, 4C | 4.47 | 0.001 | 1.98 | 6.95 |
| Temperature30:Time72 | Time0, 4C | 6.39 | <0.001 | 3.91 | 8.88 |
| TemperatureRT:NCC_2 | 4C, NCC_1 | 0.75 | 0.429 | -1.12 | 2.61 |
| Temperature30:NCC_2 | 4C, NCC_1 | 4.08 | <0.001 | 2.22 | 5.95 |

**Stratifications**

**Stability in CE + DNA extraction samples stored at 4C**

| Variable | Reference | Estimate ΔCq | P-value | Lower CI | Upper CI |
|---|---|---|---|---|---|
| Time24 | Time0 | -0.84 | 0.148 | -1.99 | 0.31 |
| Time48 | Time0 | -0.88 | 0.131 | -2.03 | 0.27 |
| Time72 | Time0 | 1.01 | 0.083 | -0.14 | 2.16 |
| Concentration10000 | Concentration1000 | -3.09 | <0.001 | -3.91 | -2.28 |
| NCC_2 | NCC_1 | 5.90 | <0.001 | 5.03 | 6.76 |

**Stability in samples CE + DNA extraction stored at RT**

| Variable | Reference | Estimate ΔCq | P-value | Lower CI | Upper CI |
|---|---|---|---|---|---|
| Time24 | Time0 | 0.54 | 0.576 | -1.40 | 2.49 |
| Time48 | Time0 | 3.44 | 0.001 | 1.50 | 5.38 |
| Time72 | Time0 | 5.48 | <0.001 | 3.54 | 7.42 |
| Concentration10000 | Concentration1000 | -3.31 | <0.001 | -4.69 | -1.94 |
| NCC_2 | NCC_1 | 6.64 | <0.001 | 5.19 | 8.10 |

**Stability in samples CE + DNA extraction stored at 30**

| Variable | Reference | Estimate ΔCq | P-value | Lower CI | Upper CI |
|---|---|---|---|---|---|
| Time24 | Time0 | 4.86 | <0.001 | 2.76 | 6.97 |
| Time48 | Time0 | 7.79 | <0.001 | 5.68 | 9.89 |
| Time72 | Time0 | 7.40 | <0.001 | 5.30 | 9.51 |
| Concentration10000 | Concentration1000 | -1.85 | 0.016 | -3.34 | -0.36 |
| NCC_2 | NCC_1 | 9.98 | <0.001 | 8.40 | 11.56 |

**Stability for NCC2 CE + DNA extraction samples at 4C**

| Variable | Reference | Estimate ΔCq | P-value | Lower CI | Upper CI |
|---|---|---|---|---|---|
| Time24 | Time0 | -0.87 | 0.263 | -2.42 | 0.69 |
| Time48 | Time0 | -0.75 | 0.333 | -2.30 | 0.81 |
| Time72 | Time0 | 1.68 | 0.035 | 0.12 | 3.24 |
| Concentration10000 | Concentration1000 | -2.94 | <0.001 | -4.04 | -1.84 |

**Stability for NCC2 CE + DNA extraction samples at RT**

| Variable | Reference | Estimate ΔCq | P-value | Lower CI | Upper CI |
|---|---|---|---|---|---|
| Time24 | Time0 | 0.62 | 0.646 | -2.10 | 3.33 |
| Time48 | Time0 | 3.92 | 0.006 | 1.20 | 6.64 |
| Time72 | Time0 | 6.70 | <0.001 | 3.98 | 9.42 |
| Concentration10000 | Concentration1000 | -3.38 | 0.001 | -5.30 | -1.46 |

**Stability for NCC2 CE + DNA extraction samples at 30**

| Variable | Reference | Estimate ΔCq | P-value | Lower CI | Upper CI |
|---|---|---|---|---|---|
| Time24 | Time0 | 6.99 | <0.001 | 4.38 | 9.60 |
| Time48 | Time0 | 10.05 | <0.001 | 7.45 | 12.66 |
| Time72 | Time0 | 9.23 | <0.001 | 6.62 | 11.83 |
| Concentration10000 | Concentration1000 | -1.38 | 0.136 | -3.22 | 0.46 |

**Stability for NCC_1 CE + DNA extraction samples at 4C**

| Variable | Reference | Estimate ΔCq | P-value | Lower CI | Upper CI |
|---|---|---|---|---|---|
| Time24 | Time0 | -0.79 | 0.306 | -2.40 | 0.83 |
| Time48 | Time0 | -1.14 | 0.148 | -2.76 | 0.47 |
| Time72 | Time0 | -0.32 | 0.667 | -1.94 | 1.29 |
| Concentration10000 | Concentration1000 | -3.39 | <0.001 | -4.53 | -2.24 |

**Stability for NCC_1 CE + DNA extraction samples at RT**

| Variable | Reference | Estimate ΔCq | P-value | Lower CI | Upper CI |
|---|---|---|---|---|---|
| **Time24** | Time0 | 0.40 | 0.697 | -1.78 | 2.58 |
| **Time48** | Time0 | 2.48 | 0.029 | 0.30 | 4.66 |
| **Time72** | Time0 | 3.04 | 0.011 | 0.86 | 5.22 |
| **Concentration10000** | Concentration1000 | -3.18 | 0.001 | -4.72 | -1.64 |

**Stability for NCC_1 CE + DNA extraction samples at 30**

| Variable | Reference | Estimate ΔCq | P-value | Lower CI | Upper CI |
|---|---|---|---|---|---|
| Time24 | Time0 | 0.61 | 0.480 | -1.22 | 2.44 |
| Time48 | Time0 | 3.25 | 0.002 | 1.42 | 5.08 |
| Time72 | Time0 | 3.76 | 0.001 | 1.93 | 5.59 |
| Concentration10000 | Concentration1000 | -2.78 | 0.001 | -4.08 | -1.49 |

**CE-extraction-free stability over temperature and time**

| **Full model** | **Reference** | **Estimate ΔCq** | **P-value** | **Lower CI** | **Upper CI** |
|---|---|---|---|---|---|
| TemperatureRT | 4C | -0.91 | 0.265 | -2.53 | 0.70 |
| Temperature30 | 4C | -3.27 | <0.001 | -4.88 | -1.66 |
| Time24 | Time0 | 0.31 | 0.641 | -1.01 | 1.63 |
| Time48 | Time0 | 0.67 | 0.315 | -0.65 | 1.99 |
| Time72 | Time0 | 0.50 | 0.452 | -0.81 | 1.82 |
| Concentration10000 | Concentration1000 | -3.28 | <0.001 | -3.82 | -2.74 |
| NCC_2 | NCC_1 | -0.38 | 0.453 | -1.36 | 0.61 |
| **Interactions** | | | | | |
| TemperatureRT:Time24 | Time0, 4C | -1.45 | 0.125 | -3.32 | 0.41 |
| Temperature30:Time24 | Time0, 4C | 0.63 | 0.504 | -1.23 | 2.49 |
| TemperatureRT:Time48 | Time0, 4C | -0.29 | 0.761 | -2.15 | 1.58 |
| Temperature30:Time48 | Time0, 4C | 4.24 | <0.001 | 2.38 | 6.10 |
| TemperatureRT:Time72 | Time0, 4C | 0.73 | 0.437 | -1.13 | 2.60 |
| Temperature30:Time72 | Time0, 4C | 5.80 | <0.001 | 3.94 | 7.67 |
| TemperatureRT:NCC_2 | 4C, NCC_1 | 1.37 | 0.055 | -0.03 | 2.77 |
| Temperature30:NCC_2 | 4C, NCC_1 | 4.90 | <0.001 | 3.51 | 6.30 |

**Stratifications**

**Stability in CE-extraction-free samples stored at 4C**

| **Variable** | **Reference** | **Estimate ΔCq** | **P-value** | **Lower CI** | **Upper CI** |
|---|---|---|---|---|---|
| Time24 | Time0 | 0.31 | 0.340 | -0.34 | 0.96 |
| Time48 | Time0 | 0.67 | 0.043 | 0.02 | 1.32 |
| Time72 | Time0 | 0.50 | 0.127 | -0.15 | 1.15 |
| Concentration10000 | Concentration1000 | -3.21 | <0.001 | -3.67 | -2.75 |
| NCC_2 | NCC_1 | -0.38 | 0.127 | -0.86 | 0.11 |

**Stability in CE-extraction-free samples stored at RT**

| **Variable** | **Reference** | **Estimate ΔCq** | **P-value** | **Lower CI** | **Upper CI** |
|---|---|---|---|---|---|
| Time24 | Time0 | -1.14 | 0.017 | -2.07 | -0.21 |
| Time48 | Time0 | 0.39 | 0.406 | -0.54 | 1.31 |
| Time72 | Time0 | 1.24 | 0.010 | 0.31 | 2.16 |
| Concentration10000 | Concentration1000 | -3.36 | <0.001 | -4.02 | -2.71 |
| NCC_2 | NCC_1 | 0.99 | 0.006 | 0.30 | 1.69 |

**Stability in CE-extraction-free samples stored at 30**

| **Variable** | **Reference** | **Estimate ΔCq** | **P-value** | **Lower CI** | **Upper CI** |
|---|---|---|---|---|---|
| Time24 | Time0 | 0.94 | 0.360 | -1.11 | 2.99 |
| Time48 | Time0 | 4.91 | <0.001 | 2.86 | 6.97 |
| Time72 | Time0 | 6.31 | <0.001 | 4.25 | 8.36 |
| Concentration10000 | Concentration1000 | -3.27 | <0.001 | -4.72 | -1.82 |
| NCC_2 | NCC_1 | 4.53 | <0.001 | 2.99 | 6.07 |

**Stability for NCC2 CE-extraction-free samples at 4C**

| Variable | Reference | Estimate ΔCq | P-value | Lower CI | Upper CI |
|---|---|---|---|---|---|
| Time24 | Time0 | 0.40 | 0.329 | -0.42 | 1.22 |
| Time48 | Time0 | 0.45 | 0.272 | -0.37 | 1.27 |
| Time72 | Time0 | 0.53 | 0.195 | -0.29 | 1.35 |
| Concentration10000 | Concentration1000 | -3.32 | <0.001 | -3.90 | -2.74 |

**Stability for NCC2 CE-extraction-free samples at RT**

| Variable | Reference | Estimate ΔCq | P-value | Lower CI | Upper CI |
|---|---|---|---|---|---|
| Time24 | Time0 | -0.57 | 0.311 | -1.71 | 0.56 |
| Time48 | Time0 | 0.77 | 0.178 | -0.37 | 1.90 |
| Time72 | Time0 | 2.00 | 0.001 | 0.87 | 3.14 |
| Concentration10000 | Concentration1000 | -3.58 | <0.001 | -4.39 | -2.78 |

**Stability for NCC2 CE-extraction-free samples at 30**

| Variable | Reference | Estimate ΔCq | P-value | Lower CI | Upper CI |
|---|---|---|---|---|---|
| Time24 | Time0 | 2.88 | 0.027 | 0.35 | 5.40 |
| Time48 | Time0 | 7.21 | <0.001 | 4.69 | 9.74 |
| Time72 | Time0 | 8.50 | <0.001 | 5.97 | 11.02 |
| Concentration10000 | Concentration1000 | -3.08 | 0.001 | -4.86 | -1.29 |

**Stability for NCC_1 CE-extraction-free samples at 4C**

| Variable | Reference | Estimate ΔCq | P-value | Lower CI | Upper CI |
|---|---|---|---|---|---|
| Time24 | Time0 | 0.14 | 0.814 | -1.13 | 1.41 |
| Time48 | Time0 | 1.12 | 0.078 | -0.15 | 2.39 |
| Time72 | Time0 | 0.45 | 0.455 | -0.82 | 1.71 |
| Concentration10000 | Concentration1000 | -2.99 | <0.001 | -3.88 | -2.09 |

**Stability for NCC_1 CE-extraction-free samples at RT**

| Variable | Reference | Estimate ΔCq | P-value | Lower CI | Upper CI |
|---|---|---|---|---|---|
| **Time24** | Time0 | -2.28 | 0.009 | -3.85 | -0.71 |
| **Time48** | Time0 | -0.38 | 0.610 | -1.95 | 1.20 |
| **Time72** | Time0 | -0.30 | 0.687 | -1.87 | 1.28 |
| **Concentration10000** | Concentration1000 | -2.92 | <0.001 | -4.03 | -1.80 |

**Stability for NCC_1 CE-extraction-free samples at 30**

| Variable | Reference | Estimate ΔCq | P-value | Lower CI | Upper CI |
|---|---|---|---|---|---|
| Time24 | Time0 | -2.93 | 0.002 | -4.55 | -1.30 |
| Time48 | Time0 | 0.32 | 0.678 | -1.31 | 1.94 |
| Time72 | Time0 | 1.92 | 0.025 | 0.29 | 3.54 |
| Concentration10000 | Concentration1000 | -3.65 | <0.001 | -4.80 | -2.50 |

**CE stability over freeze thaw**

| Full model | Reference | Estimate ΔCq | P-value | Lower CI | Upper CI |
|---|---|---|---|---|---|
| Temperature-20 | -80C | 0.00 | 1.000 | -1.96 | 1.96 |
| ft_cycle1 | ft_cycle0 | 0.58 | 0.559 | -1.38 | 2.55 |
| ft_cycle2 | ft_cycle0 | 1.64 | 0.101 | -0.32 | 3.61 |
| ft_cycle3 | ft_cycle0 | 2.34 | 0.020 | 0.37 | 4.30 |
| Storageraw | gly | 4.50 | <0.001 | 3.52 | 5.48 |
| Concentration10000 | Concentration1000 | -3.50 | <0.001 | -4.48 | -2.52 |
| NCC_2 | NCC_1 | 4.21 | <0.001 | 3.17 | 5.25 |
| **Interactions** | | | | | |
| ft_cycle1:NCC_2 | ft_cycle0, NCC1 | 1.76 | 0.272 | -1.39 | 4.90 |
| ft_cycle2:NCC_2 | ft_cycle0, NCC1 | 0.81 | 0.584 | -2.12 | 3.75 |
| ft_cycle3:NCC_2 | ft_cycle0, NCC1 | 0.42 | 0.778 | -2.51 | 3.35 |
| Storageraw:ft_cycle1 | ft_cycle0, gly | -0.77 | 0.588 | -3.55 | 2.02 |
| Storageraw:ft_cycle2 | ft_cycle0, gly | -0.81 | 0.567 | -3.59 | 1.97 |
| Storageraw:ft_cycle3 | ft_cycle0, gly | 1.62 | 0.251 | -1.16 | 4.41 |

**Stability in CE samples stored at -80**

| Full model | Reference | Estimate ΔCq | P-value | Lower CI | Upper CI |
|---|---|---|---|---|---|
| Storageraw | gly | 3.79 | <0.001 | 2.58 | 5.00 |
| ft_cycle1 | ft_cycle0 | 0.58 | 0.501 | -1.13 | 2.30 |
| ft_cycle2 | ft_cycle0 | 1.64 | 0.060 | -0.07 | 3.36 |
| ft_cycle3 | ft_cycle0 | 2.34 | 0.008 | 0.62 | 4.05 |
| Concentration10000 | Concentration1000 | -3.16 | <0.001 | -4.38 | -1.95 |
| NCC_2 | NCC_1 | 4.38 | <0.001 | 3.09 | 5.66 |

**Stability in CE samples stored at -20**

| Full model | Reference | Estimate ΔCq | P-value | Lower CI | Upper CI |
|---|---|---|---|---|---|
| Storageraw | gly | 5.21 | <0.001 | 3.65 | 6.78 |
| ft_cycle1 | ft_cycle0 | 1.53 | 0.174 | -0.69 | 3.74 |
| ft_cycle2 | ft_cycle0 | 4.46 | <0.001 | 2.25 | 6.68 |
| ft_cycle3 | ft_cycle0 | 4.39 | <0.001 | 2.17 | 6.60 |
| Concentration10000 | Concentration1000 | -3.83 | <0.001 | -5.40 | -2.26 |
| NCC_22 | NCC_1 | 4.04 | <0.001 | 2.38 | 5.70 |

**SD stability over freeze thaw**

| Full model | Reference | Estimate ΔCq | P-value | Lower CI | Upper CI |
|---|---|---|---|---|---|
| Temperature-20 | -80C | 0.12 | 0.507 | -0.23 | 0.47 |
| ft_cycle1 | ft_cycle0 | -0.16 | 0.500 | -0.64 | 0.32 |
| ft_cycle2 | ft_cycle0 | -0.03 | 0.921 | -0.53 | 0.48 |
| ft_cycle3 | ft_cycle0 | 0.47 | 0.058 | -0.02 | 0.95 |
| Storageraw | gly | -2.22 | <0.001 | -2.57 | -1.87 |
| Concentration10000 | Concentration1000 | -3.02 | <0.001 | -3.36 | -2.67 |
| NCC_2 | NCC_1 | -0.14 | 0.470 | -0.51 | 0.23 |
